# MetaboShiny – interactive processing, analysis and annotation of direct infusion metabolomics data

**DOI:** 10.1101/734236

**Authors:** Joanna C. Wolthuis, Stefania Magnusdottir, Mia Pras-Raves, Maryam Moshiri, Judith J.M. Jans, Boudewijn Burgering, Saskia van Mil, Jeroen de Ridder

**Author notes:** **Correspondence should be addressed to:** Jeroen de Ridder, or Saskia van Mil, UMC Utrecht, Center for Molecular Medicine, STR3.217, PO Box 85060, 3508 AB Utrecht, The Netherlands. Joint senior authors (Saskia van Mil, Jeroen de Ridder).

## Abstract

Direct infusion untargeted metabolomics, as mass-over-charge values and intensity of ions, allows for rapid insight into a sample’s metabolic activity. However, analysis is often complicated by the large array of detected m/z values and the difficulty to prioritize important m/z and simultaneously annotate their putative identities. To address this challenge, we developed MetaboShiny, a novel R/RShiny-based metabolomics package featuring data analysis, database- and formula-prediction-based annotation and visualization. To demonstrate this, we reproduce and further explore a MetaboLights metabolomics bioinformatics study on lung cancer patient urine samples. MetaboShiny enables rapid and rigorous analysis and interpretation of direct infusion untargeted metabolomics data.

## Introduction

Metabolomics is the underlying biochemical layer of the genome, transcriptome and proteome, which reflects all the information expressed and modulated by these omics layers. Because metabolomics provides an almost direct readout of metabolic activity in the organism, metabolomics can be used to diagnose diseases from biofluids, discover new drugs and drug targets, and further precision medicine(Wishart, 2016).

A common method to acquire metabolomics data is mass spectrometry (MS), which records the input metabolites’ mass to charge ratios (m/z). An example of a metabolomics method is direct infusion mass spectrometry (DI-MS), which detects tens to hundreds of thousands of m/z values representing metabolites at single part per million (ppm) accuracy (de Sain-van der Velden *et al.*, 2017). DI-MS runtimes are in the order of one minute per sample, making it highly suitable for high-throughput applications, such as for instance in diagnostics applications. Problematically, DI-MS routinely produces over a hundred thousand unidentified m/z values, which makes user interpretation of the data exceedingly challenging (Lin *et al.*, 2010; Schrimpe-Rutledge *et al.*, 2016; de Sain-van der Velden *et al.*, 2017).

Non-direct MS methods like liquid/gas-chromatography mass spectrometry (LC/GC-MS) and fragmentation mass spectrometry (MS/MS) generally involve *a priori* metabolite selection based on a chosen compound feature, e.g. polarity or, in MS/MS, precursor m/z value, while DI-MS metabolomics is performed on the input sample without pre-filtering the sample. Before this data can be effectively utilized to answer diagnostic or research questions, m/z values need to be matched to a potential metabolite identity, commonly called metabolite *annotation*.

Most freely available tools use a single database for metabolite annotation, thus limiting the number of metabolite identities that can be matched in one search query. For instance, many existing tools rely on a fixed integrated version of publicly available databases, with no opportunity to update to the most recent version (Misra and Mohapatra, 2018). However, many databases provide valuable contextual information, such as the FooDB, which focuses on compiling research on compounds in food and their metabolites(www.foodb.ca). With the many databases included in MetaboShiny, users are more likely to find a database that fits their line of research and gives relevant contextual information on each compound.

Moreover, annotation is only one step in the data analysis workflow and needs to be integrated into downstream analytics, such as data normalization, statistical testing, visualization, clustering or supervised machine learning. Many of the currently available tools are only geared towards a subset of these tasks, impeding ease-of-use and time-to-results.

Given the potential of DI-MS data, we sought to address these issues and enhance the strengths of the available tools and packages. The resulting software package, MetaboShiny, supports the user in performing a range of common MS data analysis steps such as normalization, statistical analysis and putative annotation of m/z values. MetaboShiny is built in the interactive *Shiny* framework for the R programming language (Chang *et al.*, 2015). MetaboShiny, through our included companion package *MetaDBparse*, supports over 30 databases (Table S1) and is capable of searching through millions of m/z values, including metabolite variants with over twenty commonly observed adducts and isotopes (*Table S2, S3*). Additionally, users can create their own user-defined databases and annotation and statistical analysis can be performed in parallel, allowing the user to quickly find putative identities for m/z values.

MetaboShiny supports annotation through matching the highly accurate m/z value to a database or deducing a molecular formula based on organic chemistry and pre-defined rules such as the *Seven Golden Rules* that take into account ratios of oxygen, nitrogen and hydrogen atoms (Kind and Fiehn, 2007; Schrimpe-Rutledge *et al.*, 2016). MetaboShiny performs downstream data analysis directly on the m/z ratios and metabolite annotation occurs only on e.g. significant hits that result from the analysis. Installation is facilitated by Docker (Anderson, 2015), which pre-installs all required libraries and packages with a single button click.

## Methods

MetaboShiny is an R based application, wrapped in a ‘*Shiny’* user interface(Chang *et al.*, 2015). It enables users to fluidly switch between running statistical analyses, exploring plots, tables and m/z annotation (*Figure 1*). Furthermore, by summarizing the results in publication-ready figures, which can be exported in multiple formats including vector formats, MetaboShiny decreases the amount of time needed to get from initial data to insights. A flowchart of the underlying processes is provided in *Figure S1*. Additionally, a comparison with other freely available metabolomics software tools can be found in *Section S3* and *Table S1*.

**Figure 1:**
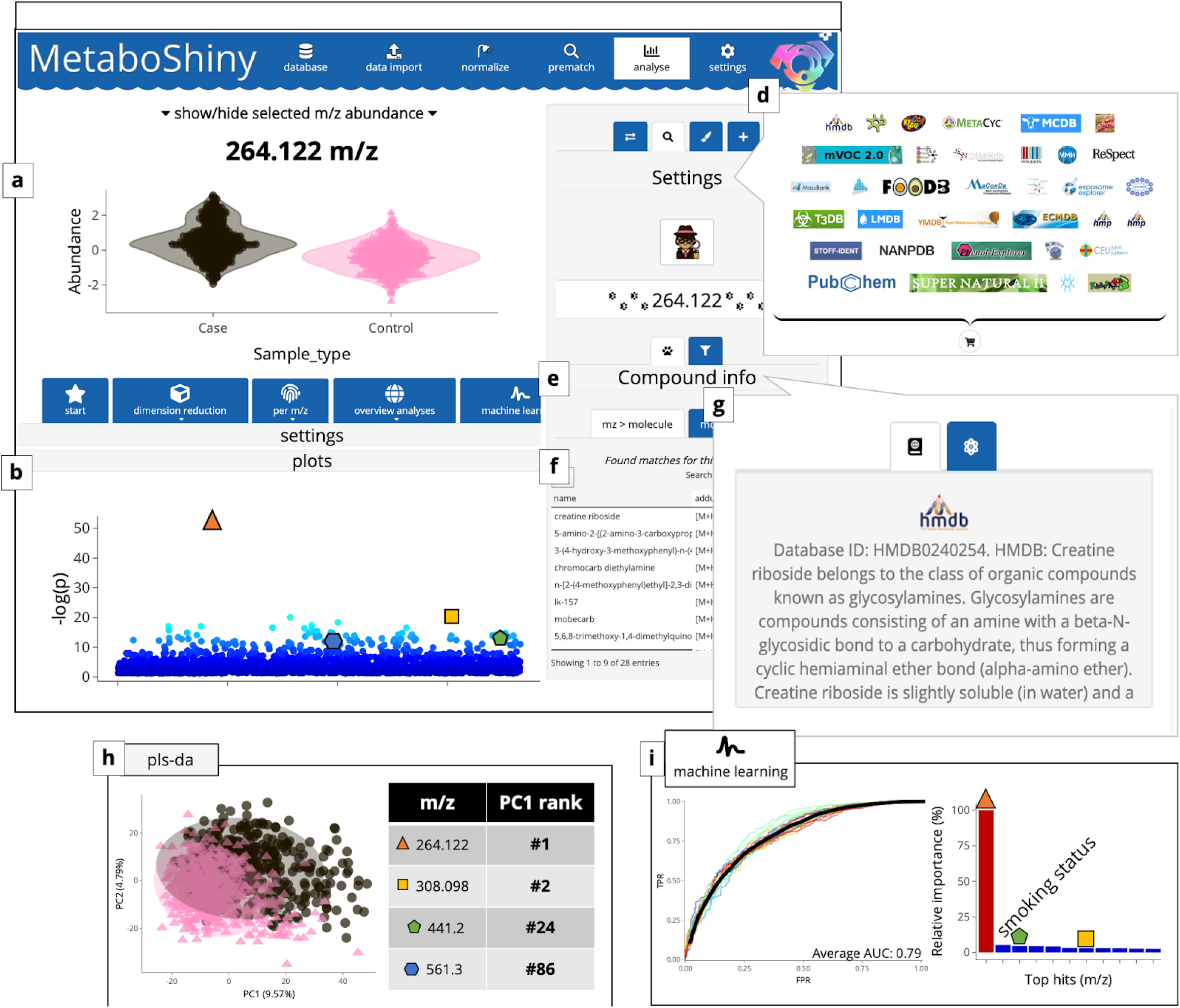
Overview of the MetaboShiny application. The displayed results themselves are obtained from a MetaboLights dataset MTBLS28, which is more extensively discussed in section S1. (a) Box plot of the abundance of a single metabolite. Aside from beeswarm- and boxplots, scatter/violin plots are available. (b) Manhattan-like plot of all t-test hits. (c) Extra options for the searching algorithm. (d) Subset of databases available for search functionality. Users can select the ones they prefer. (e-g) The search results section of the sidebar. Users can scroll through the search results and display the database description and structure of the metabolite selected in that table. Results were filtered first by the main peak isotopes, and subsequently sorted by ppm error. Compounds with identical molecular formulas, but different structures/SMILES are listed as separate rows. Compounds with identical SMILES but different names (IUPAC, commercial, etc) are collapsed into one row. Users can then view the synonyms and descriptions once they access the detail view for the selected search hit. (h) PLS-DA plot and loading results. (i) Examples of machine learning results. Users see ROC curves for one or multiple models (and their average performance) and an overview of variable importance (including metadata if the user wishes).

### Metadata

MetaboShiny supports the MetaboLights metadata format in order to maximize compatibility with existing metabolomics data analysis software. Moreover, MetaboShiny includes a novel meta-data format which is in line with state-of-the-art data stewardship best practises in order to better adhere to the FAIR principles(Rothfritz, 2019). Examples of metadata tables and peak tables compatible with MetaboShiny are given in *table S6* and *S7* for metadata and peak data respectively.

### Statistics

MetaboShiny offers multiple modes of statistical analysis, with various options for bivariate, multivariate, time series and two-factor experimental variables using paired or unpaired samples. Through the integrated *MetaboAnalystR* package, users have access to t-tests, fold-change analysis, pattern analysis, various dimension reduction methods such as PCA and PLS-DA, and volcano plots (Chong *et al.*, 2018). Additionally, users have access to t-SNE, heatmaps and machine learning. We demonstrate examples of these analyses in the interface in figure 1, where the t-test (*figure 1a-b)*, PLS-DA (*figure 1h)* and random forest machine learning (*figure 1j)* are used as examples. This example data is further discussed in *section S1* of the Supplemental Materials.

### Subsetting, switching and intersecting

MetaboShiny offers to explore specific subsets of the data or select another experimental variable from supplied metadata. Results from all analyses and subsets may be compared through Venn diagrams, which offer hypergeometric testing to test the significance of overlapping hits and are useful for prioritizing multiple significant m/z values (*figure S3a)*.

### Machine learning

*MetaboAnalystR* natively supports several machine learning methods, MetaboShiny moreover leverages the *caret* package (Kuhn, 2008), which is currently the most extensive machine learning package available for R. Flexibility is maximized by allowing users to create predictive models from almost a hundred different methods with manually tunable parameters. The available methods have been selected based on their ability to provide variable importance scores. Furthermore, some models are not designed for multivariate data. Using these considerations, MetaboShiny offers caret models based on the dataset the user is working on. Users can include metadata in their predictive models alongside m/z values, and explore model variable importance for m/z prioritization (*figure 1i)*.

### Interactivity

An important and unique feature of MetaboShiny is that it offers data interactivity through the *DT* and *plotly* R packages (Sievert *et al.*, 2016; Xie, 2017), allowing for clicking on and magnification of specific points or regions of interest in plots or tables that may be filtered and sorted, and immediately continuing to the m/z value annotation step.

### Annotation

Currently available m/z annotation methods are limited by searches in a single or limited set of databases because searching all available databases is very time consuming. For this reason, MetaboShiny streamlines the process of m/z annotation, enabling the user to rapidly retrieve matches in a wide range of optimized compound databases that include adducts and isotopes, according to a predetermined error margin (*Figure S1a)*. Before searching for a molecular identity of an m/z value, users select the databases of interest. MetaboShiny currently supports 35 compound databases (see *Table S1* for an overview of all database options). Our *MetaDBparse* package subsequently generates m/z values for common isotopes and adducts known to form for each compound in the database (*Figure S1a, Table S2-S3*). Compound databases are generated and stored locally by the user. Additionally, a subset of online-only databases are available. MetaboShiny moreover allows users to interactively explore and filter the retrieved annotations through interactive summary figures. According to the Metabolomics Standards Initiative, MetaboShiny provinces MSI level 2 annotation i.e. putatively annotated compounds (Members: and MSI Board Members:, 2007). It should be noted, that for definitive identification of m/z values, validation using authentic chemical standards in the same laboratory is required.

### Availability

MetaboShiny is available on Docker Hub under the *jcwolthuis/MetaboShiny* repository. The source code is available on GitHub in the *joannawolthuis/MetaboShiny* repository. If building from source, administrator rights are needed to install the necessary libraries. The *MetaDBparse* package containing all the search and compound-db specific functionality without the *Shiny* user interface is available from the *joannawolthuis/MetaDBparse* repository. For ease of use, Docker is recommended to avoid the complications arising from installing dependencies. The user manual is also available on the GitHub repository for *MetaboShiny*.

## Results and discussion

### Functionality of MetaboShiny demonstrated using a test dataset

To demonstrate the functionality of MetaboShiny, we leveraged the LC-MS dataset on urine samples from 1005 patients with and without lung cancer. This work identified 4 m/z values at MSI level 1, *264 m/z (* Δ in figures), *308 m/z* (□), *441 m/z* (□) and *561 m/z* (○) predictive for cancer status, which were confirmed through targeted mass spectrometry, one of which, namely 264 m/z, was a novel compound called *creatine riboside (Members: and MSI Board Members:, 2007; Mathé et al., 2014)*..

As an initial analysis, we ran a t-test in MetaboShiny(*figure 1a-b)* and this returned a list of most significantly different m/z values between the control and disease groups. When using MetaboShiny to search for potential annotations for the m/z value at the top of this list, a search including the HMDB (*figure 1c-d)* uncovered creatine riboside (M+H adduct) as a putative hit (*figure 1f)*. The built-in HMDB compound description page (*figure 1e)* reports that this m/z feature was first discovered by Mathé *et al(Mathé et al., 2014)* (*figure 1g*), confirming the validity of this finding. All this information can be retrieved with a few mouse clicks.

Additionally, PLS-DA analysis was explored. This yielded a model that achieved significant separation between the population and lung cancer groups (p < 0.03; *figure 1h*). The loading for PC1 contains the four m/z values of interest, ranked #1, #2, #24 and #86 respectively. These results indicate that the four compounds are not just significant in univariate t-tests, but also can be used to train predictive models that stratify samples by cancer status.

MetaboShiny can also be used to perform more routine analyses such as fold-change analysis, volcano plots and correlation analyses. Furthermore, heatmaps and venn diagrams are available.

Lastly, Mathé et al. trained a random forest model and combined the results of multiple classifiers to find compounds that were good predictors in multiple metadata groups (sex, race, and smoking status)(Mathé *et al.*, 2014). Using MetaboShiny this analysis can be completely reproduced within an hour, demonstrating the utility of the software for rapid hypothesis generation and biomarker discovery.

Two of the four metabolites that Mathé et al. identified rank within the top 20 (#1 and #3 for 264 and 308 m/z respectively) P7L9 in terms of variable importance, with *264 m/z* having the highest predictive value. Aside from m/z values, smoking status was a strong predictor for disease (ranked #2 by the Random Forest, *figure 1j*). Additional analyses exploring further functionality such as subsetting, formula prediction and PubMed text mining are available in *Supplemental Section S2*.

Taken together, this demonstrates the power of MetaboShiny to achieve rapid hypothesis generation, which can be followed up with additional experiments.

### Performance quantification of MetaboShiny

To quantify MetaboShiny’s speed, a quantitative analysis was performed on MetaboShiny’s database searching performance. For an informative comparison, we created 300 databases spanning the range of real world database sizes. For each of these database sizes, a random pool of 100 m/z values, evenly distributed over the 60 – 600 m/z range were used individually to perform a match search.

The machine used was a MacBook Pro (13-inch, 2016) with 16GB RAM and a 2,9 GHz Intel Core i5 processor. M/z values with more matches in the database trend towards taking more time, as MetaboShiny fetches additional information (description, molecular formula, name) when a match is found. Furthermore, SQLITE occasionally performs self-maintenance(indexing, for example), or the test system may simultaneously be performing background tasks. This is likely to cause some of the slower outliers. Regardless of this, one search even at the current maximum database size does not exceed 0.8 seconds for a single search, generally staying below half a second(*Figure 2a)*.

**Figure 2:**
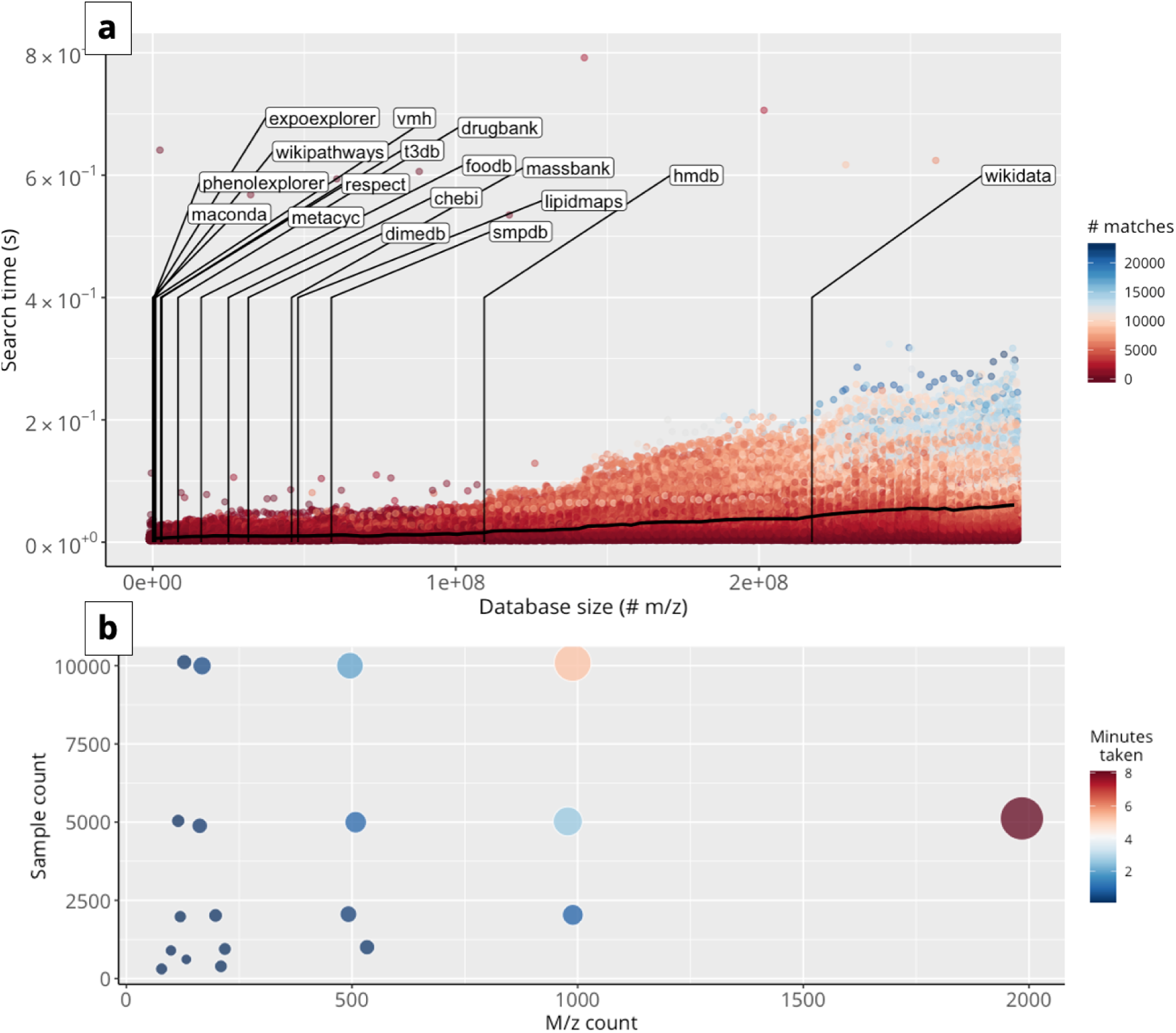
Analysis of MetaboShiny’s speed performance. (a) Time to annotate a m/z value. Searching one m/z value takes longer if more matches are found. On average, even with large databases including 3e8 m/z values, performing a search on a single m/z value takes under one second. Labels show the size of various included databases with Wikidata being the most extensive. Color of points represents the amount of matches found for each m/z value. (b) Time to process data from import to end of normalization. Size and color of markers represents minutes.

Furthermore, we tested the time it takes to import and normalize a dataset, after which analysis can be started. The test dataset took five minutes to process to the point of being able to start analysis. We generated synthetic datasets consisting of combinations of a minimum of 10 and a maximum of 10000 samples, and minimally 10 to maximally 2000 m/z values. Importing and normalization generally takes less than 10 minutes, this time increasing as more m/z values and/or samples are added in the dataset. This was using the same normalization settings as the test dataset. Missing values were imputed using Random Forest, other normalization methods are expected to run substantially faster (*Figure 2b*).

### Post-annotation considerations

Putative identities of m/z values will still require validation through LC/MS-MS. MetaboShiny guides users to this step by prioritizing compounds based on their isotope and adduct status, alongside using database compound descriptions to prioritize compounds based on what is biologically known. (Coley *et al.*, 2019).

Further plans to enhance MetaboShiny include refining *in silico* compound annotation and prioritization, expanding the amount of available databases, and increasing the ability to streamline data integration.

## Acknowledgements

We thank Arie Kies and Pim Langhout (DSM) for critical discussions, Daphne van Beek and Jasmin Böhmer for assistance with data stewardship, and Marc Pages Gallego for testing and input on the software.

## Compliance with Ethical Standards

### Funding contributions

This research is supported by the Dutch Technology Foundation STW, which is the Applied Science Division of NWO, and Technology Programme of the Ministry of Economic Affairs. This research is also supported by DSM Nutritional Products. JdR is supported by a Vidi Fellowship (639.072.715) from the Dutch Organization for Scientific Research (Nederlandse Organisatie voor Wetenschappelijk Onderzoek, NWO)

### Software

R 3.5.1, SQLITE, Docker – exact packages detailed in featured package info.

### Data

The featured data was previously uploaded to MetaboLights, using the identifier “*MTBLS28*”. For information on how this data was collected, please refer to their manuscript.

## References

Anderson, C. (2015) ‘Docker [Software engineering]’, IEEE Software, 32(3), pp. 102–c3. doi: 10.1109/MS.2015.62.

Chang, W. et al. (2015) ‘Shiny: web application framework for R’, R package version 0. 11, 1(4), p. 106.

Chong, J. et al. (2018) ‘MetaboAnalyst 4.0: towards more transparent and integrative metabolomics analysis’, Nucleic acids research, 46(W1), pp. W486–W494. doi: 10.1093/nar/gky310.

Coley, C. W. et al. (2019) ‘A graph-convolutional neural network model for the prediction of chemical reactivity’, Chemical science, 10(2), pp. 370–377. doi: 10.1039/c8sc04228d.

Kind, T. and Fiehn, O. (2007) ‘Seven Golden Rules for heuristic filtering of molecular formulas obtained by accurate mass spectrometry’, BMC bioinformatics, 8, p. 105. doi: 10.1186/1471-2105-8-105.

Kuhn, M. (2008) ‘Building predictive models in R using the caret package’, Journal of statistical software. math.chalmers.se. Available at: http://www.math.chalmers.se/Stat/Grundutb/GU/MSA220/S18/caret-JSS.pdf.

Lin, L. et al. (2010) ‘Direct infusion mass spectrometry or liquid chromatography mass spectrometry for human metabonomics? A serum metabonomic study of kidney cancer’, The Analyst, 135(11), pp. 2970–2978. doi: 10.1039/c0an00265h.

Mathé, E. A. et al. (2014) ‘Noninvasive urinary metabolomic profiling identifies diagnostic and prognostic markers in lung cancer’, Cancer research, 74(12), pp. 3259–3270. doi: 10.1158/0008-5472.CAN-14-0109.

Members:, M. B. and MSI Board Members: (2007) ‘The Metabolomics Standards Initiative’, Nature Biotechnology, pp. 846–848. doi: 10.1038/nbt0807-846b.

Misra, B. B. and Mohapatra, S. (2018) ‘Tools and resources for metabolomics research community: A 2017–2018 update’, Electrophoresis. Wiley Online Library. Available at: https://onlinelibrary.wiley.com/doi/abs/10.1002/elps.201800428.

Rothfritz, L. (2019) ‘The FAIR data principles’. doi: 10.14293/s2199-1006.1.sor-compsci.clnbrup.v1.

de Sain-van der Velden, M. G. M. et al. (2017) ‘Quantification of metabolites in dried blood spots by direct infusion high resolution mass spectrometry’, Analytica chimica acta, 979, pp. 45–50. doi: 10.1016/j.aca.2017.04.038.

Schrimpe-Rutledge, A. C. et al. (2016) ‘Untargeted Metabolomics Strategies-Challenges and Emerging Directions’, Journal of the American Society for Mass Spectrometry, 27(12), pp. 1897–1905. doi: 10.1007/s13361-016-1469-y.

Sievert, C. et al. (2016) ‘plotly: Create Interactive Web Graphics via “plotly. js”‘, R package version.

Wishart, D. S. (2016) ‘Emerging applications of metabolomics in drug discovery and precision medicine’, Nature reviews. Drug discovery, 15(7), pp. 473–484. doi: 10.1038/nrd.2016.32.

Xie, Y. (2017) ‘DT: a wrapper of the JavaScript library “DataTables”. 2016’, R package version 0. 2.

