## Supplemental Materials for "MetaboShiny – interactive processing, analysis and annotation of direct infusion metabolomics data"

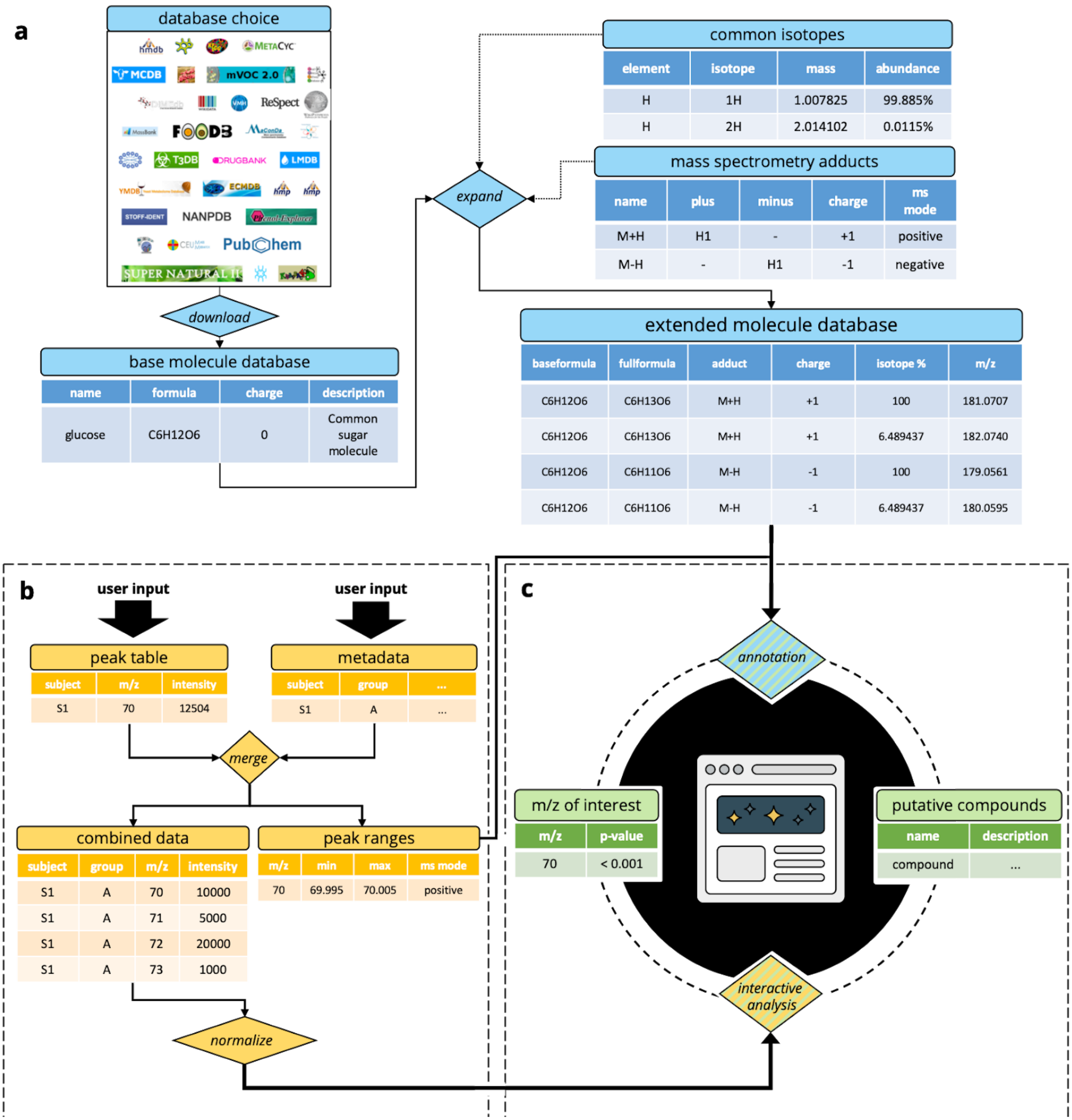


**Figure S1:** General workflow of MetaboShiny. Users trigger molecule database creation n (a), combine their peak data and metadata in (b) and enter the cycle of biomarker discovery and annotation in (c).

### Section S1. Extended methods

### Databases

When the user clicks the 'build database' button, the most recent release of the database is either downloaded from the creator or through querying the creator's web application programming interface (API). This functionality is provided by the built-in parsers of the authors' *MetaDBparse* R package. Each database only needs to be downloaded once but can be updated by re-downloading the latest version of the database. The databases contain the name, molecular formula, charge, description, and if available, structure in SMILES format(Anderson, Veith and Weininger, 1987). Currently, 45 adduct variants, 21 in positive ion mode and 24 in negative ion mode, are calculated for each molecular formula and its isotopes (*Table S2*). Users can add and remove adducts through the settings panel.

In terms of adduct creation, the structure in SMILES can be used to pre-filter which compounds can form which adducts. Users can supply a SMARTS string that finds possible chemical sites needed for the adduct to be created, also called adduct rules(Daylight Chemical Information, no date; Anderson, Veith and Weininger, 1987). If such a rule is not supplied, or no SMILES are available for a compound, the structure is not checked and adduct possibility is based on molecular formula only (presence and absence of necessary atoms for the reaction). Currently, 35 databases are available offline: the HMDB, ChEBI, DimeDB, KEGG, MetaCyc, WikiPathways, SMPDB, Wikidata, VMH, ReSpect, MassBank, MetaboLights, FooDB, MaConDa, Blood Exposome Database, LipidMaps, Exposome Explorer, Toxin and Toxin Target Database(T3DB), DrugBank, Phenol-Explorer, PAMDB, mVOC, RMDB, BMDB, STOFF-IDENT and NANPDB(O'Shea *et al.*, no date; Kanehisa and Goto, 2000; Subramaniam and Fahy, 2007; Degtyarenko *et al.*, 2008; Horai *et al.*, 2010; Guo *et al.*, 2012; Sawada *et al.*, 2012; Weber *et al.*, 2012; Rothwell *et al.*, 2013; Jewison *et al.*, 2014; Wishart *et al.*, 2015; Kale *et al.*, 2016; Ntie-Kang *et al.*, 2017; Huang *et al.*, 2018; Lemfack *et al.*, 2018; Slenter *et al.*, 2018; Wishart, Yannick D. Feunang, *et al.*, 2018; Wishart, Yannick Djoumbou Feunang, *et al.*, 2018; Barupal and Fiehn, 2019; Noronha *et al.*, 2019; Turki *et al.*, 2019; Caspi *et al.*, 2020; Neveu *et al.*, 2020).

If the user is connected to the internet, additional databases are available through two distinct methods - users can predict a chemical formula for a compound and search that formula through PubChem, ChemSpider, SUPER NATURAL II and/or KNaPsack (Dunkel *et al.*, 2006; Ayers, 2012; Nakamura *et al.*, 2014; Ihlenfeldt, 2018),. Furthermore, users can indirectly search METLIN and MASS through CEU Mass Mediator(Gil-de-la-Fuente *et al.*, 2019). Most databases are immediately available, with some exceptions - DrugBank and MetaCyc require that users create an account and download selected files first, after which MetaboShiny handles the parsing of those database files. Searching ChemSpider requires an API token, which can be input in the settings menu of MetaboShiny.

Aside from the 'base' database, MetaboShiny builds an extended database which includes adduct rule checking, adduct formula generation and isotope pattern generation to get all possible m/z values connected to the base compound. The time it takes to set up databases depends on the size and number of the databases chosen. To build them all from nothing, including isotope pattern and adduct generation, takes approximately 24 hours on our systems. This process can be sped up on systems with high computational power by using the multithreading functionality accessible in the settings menu of MetaboShiny. Updating a database takes less time once the full database is set up, as MetaDBparse only processes new SMILES (SMILES are first processed by the rcdk package and then re-output into Canonical SMILES).

#### *Metadata*

When creating a MetaboShiny project, users upload their peak table (in either .CSV or .SQLITE format) and metadata (in .XLS(X) or .CSV format). The general structure of these tables is visualized under the boxes in the directional graph in *Figure 1b*. Examples are provided in *Table S4-S5*. The metadata table requires a date of measurement, sample identifier, unique individual identifier and experimental group. If desired, more data can be added, including other specifics of the experiment such as supplement dosages, diet types or age. All of these features can be used later in subsetting and machine learning. The metadata is included by manipulating the formed *MetaboAnalystR* project (*mSet*) post-normalization within R and can be used to subset data.

#### *Data storage*

M/z error margins are stored in SQLITE in an RTree structure, which allows for fast range searches(Allen and Owens, 2010). MetaboShiny uses the mean m/z and parts per million (ppm) error margin to create an m/z range table (min - max m/z allowed to match), which is later used to quickly find matches in the downloaded compound databases. For every sample, the intensity of every m/z peak is stored, alongside the ionization mode it was found in (positive or negative). The ionization mode is important when matching specific adducts, because some adducts are only formed in either negative or positive mode.

The compound databases are also stored in SQLITE, where the tables have been set up to be in database normal form. The source database is downloaded from the host and is parsed into a format containing the name, description, formula, charge and structure of each compound using the included *MetaDBparse* companion package. These reformatted 'base' databases are saved separately per source database as separate files. The extended database containing all adducts and isotopes for each molecular structure has a different structure to avoid redundant calculation of adducts and isotopes - it does not use the compound descriptions and names but only hosts the information necessary to calculate m/z values, which is molecular formula, structure and charge. The base and extended databases are joined together when performing annotation to provide all the information necessary for the user. The differences in data structure between the base and extended tables are visualised in *Figure S1a*.

#### *Normalization*

*MetaboAnalystR* requires a .CSV file with a specific formatting(Chong and Xia, 2018). For the majority of the normalization options users can refer to the *MetaboAnalystR* package. MetaboShiny also implements such random forest missing value imputation as additional option(Stekhoven, 2015). Batch correction is based on the batch included in the metadata file, with or without quality control (QC) samples, using either the *ComBat* or *BatchCorrMetabolomics* packages depending on the presence of these QC samples(Johnson, Li and Rabinovic, 2007; Wehrens *et al.*, 2016). Processing the data from the initial file to the point of statistical analysis takes only 5 minutes for the most basic settings, although this time can vary if using more advanced normalization settings such as random forest missing value imputation.

#### *Visualization*

MetaboShiny mostly relies on the *ggplot2* package for data visualization. All of the commonly available color palettes and plot styles from *ggplot2* are available(Wickham, 2011). MetaboShiny features 2d and 3d scatter plots for dimension reduction, line plots (used for temporal analysis and machine learning plots), box/violin/beeswarm plots, heatmaps, pie charts and venn diagrams.

All these plots are interactive because of the *DT* and *plotly* R packages(Sievert *et al.*, 2016; Xie, 2017). These packages are instrumental in creating plots where users can click on specific points of interest or interactive tables that users can filter and sort on, and immediately continue to the m/z value annotation step. This interactivity is particularly useful when exploring heatmaps and volcano plots, where many methods use labels for the hundreds of points or rows. This reduces clutter and users can zoom in on selected areas and explore them at will (Figure S8).

#### *Machine learning*

For machine learning results, MetaboShiny offers a summary receiver operating characteristic (ROC) curve, the area under the curve (AUC) and a separate bar chart with variable importance for each m/z and metadata value. These are all interactive, and users can view the weight of each predictor once clicking on a specific ROC curve.

For the analysis discussed in this work, MetaboShiny built 10 random forest models for the analysis of the whole dataset, using parameters *mtry = 3167* and *ntree = 500*.

#### *Annotation*

Every time the user selects an m/z value from a plot or table, it is registered, and this '*current m/z value'* is displayed on the search tab on the sidebar that remains visible during statistical analysis. Once the user has selected adducts and databases, a search can be carried out. Since the m/z value of interest is linked to an m/z range table, which takes the ppm error margin into account, MetaboShiny will retrieve the information associated with all compounds that fall within this m/z range within all selected databases.

Results are displayed within a table with multiple pages in the sidebar, alongside a figure summarizing the words used in the descriptions, and two figures summarizing which adducts and which databases are represented in the results. To increase insights, users can choose to search in a user-specified amount of PubMed abstracts and find the most used words in these abstracts. They can also view all of the titles of these papers.

An additional functionality includes predicting possible chemical formulas based on m/z value and predetermined rules, utilizing some the *Seven Golden Rules(Kind and Fiehn, 2007)* (excluding the rules that use presence in existing databases) and the *rcdk* package(Guha and Cherto 2017). Using either the elements CHO, CHNO, CHNOP and CHNOPS as possible elements, possible formulas adding up to the m/z in question are generated. To do this, isotope patterns are calculated, from which the main isotope m/z is used to do further calculations. These formulas are then filtered according to the user's selected Golden Rules dictating which ratios of elements to one another are most likely. These rules were acquired from the Visual Basic script offered by the Fiehn lab(Kind and Fiehn, 2007). Users can choose whether to use these rules or not. Search functionality is provided by our MetaDBparse package which is included in the docker file.

### Section S2. Functionality of MetaboShiny demonstrated (Extended)

The data by Mathe et al. are publically available on MetaboLights(Kale *et al.*, 2016) (identifier *MTBLS28*). Please note that the original data comes from an LC-MS. We decided to use this data as the authors validated three of their four compounds of interest, allowing us to assess the quality of our annotation algorithms. We excluded the retention time to adhere to the DI-MS data format.

#### *Preprocessing*

The available peak tables for positive and negative mode offered a total of 3166 m/z values for each sample. These peak tables were reformatted to remove the retention time, which is currently not searchable in most databases and not used in MetaboShiny for compound annotation (script available in supplemental). Additionally, the metadata for each sample was reformatted to fit MetaboShiny's required format, although MetaboLights metadata format is also supported. Both files were imported into MetaboShiny. The conversion script and resulting files are available here. As per the original authors' protocol, an error margin of 25 parts per million (ppm) was used to perform matching (Mathé *et al.*, 2014).

Within MetaboShiny, with a single click after entering normalization preferences, data can be log transformed and normalized based on the sum of the total sample intensity to correct for large differences in overall acquired signal during mass spectrometry detection. We further Z-transformed the data to account for differences between variable distributions.

Missing values were imputed with random forest, as it was previously determined to be most accurate(Wei *et al.*, 2018). Any m/z values missing a signal in more than one percent of samples were excluded from further analysis. This resulted in 3141 metabolites.

#### *Random Forest*

Mathé et al. initially trained a random forest model and combined the results of multiple classifiers to find compounds that were good predictors in multiple metadata groups (sex, race, and smoking status)(Mathé *et al.*, 2014). Using MetaboShiny this analysis can be completely reproduced within only a few minutes, demonstrating the utility of the software for rapid hypothesis generation and biomarker discovery. The Random Forest classifier is amongst the 50+ machine learning algorithms included in Metaboshiny(Kuhn, 2008). The model performs better than random classification, with an area under the curve (AUC) of 0.84.

#### *In depth analysis of test dataset by subsetting and intersecting analyses with MetaboShiny*

MetaboShiny enables subsetting and analysis of intersecting features making it very easy for the user to subset and filter samples based on the supplied metadata. For example, as smoking is highly correlated with lung cancer risk, the dataset can be divided into three subsets based on smoking status.

To view the data from another angle, the data was analysed using the machine learning functionality of MetaboShiny. Machine learning models are capable of capturing multivariate variance that other univariate tests cannot. The full *caret* functionality included in MetaboShiny allows users to explore multiple machine learning models, including the random forest, LASSO, ridge regression, general linear models and many more(Kuhn, 2008).

For each of these subsets (*present, past* and *never smoker*) regularized (glmnet) predictive models were built with all m/z features. The predictive importance of each m/z value in these subsets is stored and can be used to do intersectional analysis. The Venn diagram using the top 50 hits for each of these three subsets produced by MetaboShiny is presented in *figure S2a*. The analysis revealed that the previously discussed compound creatine riboside (*264 m/z,* triangle in figures 1 and S2-3) is in the top 15 predictive m/z values for all three subsets, independent of smoking status *(Figure S2a).* This further validates its status as top significant compound as in the results from Mathe et al(Mathé *et al.*, 2014).

Subsequently, MetaboShiny was used to search for metabolites that only were predictive of cancer status in the group of patients that never or only formerly smoked. From the top 50 predictive m/z values in total, 44 are uniquely predictive for lung cancer status in this subset, implying a larger role for them in lung cancer in non-smokers.

To demonstrate how MetaboShiny enables users to identify putative compounds of interest, these 15 compounds were further explored. Browsing the available identities gave insight into which compounds are interesting for further validation in targeted analyses. As an example, one metabolite of interest, at *127 m/z,* was identified through browsing the possible m/z identities for metabolites found in humans through the HMDB. MetaboShiny features a side panel showing differential expression for each m/z selected (figure S2b).

Next, to find possible compound annotations for *127 m/z*, a full search for all downloaded databases was performed (supplemental table S1). In total this returned 122 matches. After disregarding duplicates, this m/z value matches hydroiodic acid (iodine in solution), and iodine-related compounds (Figure S2c). In this case only M-H adducts were considered, as the m/z value was registered in negative mode, and this is the most prevalent ionized form seen in this mode(de Sain-van der Velden *et al.*, 2017).

To aid the user in interpreting the results, MetaboShiny offers to do a general PubMed search in this case providing a general idea of the co-occurrence of iodine and cancer in literature. A user-specified (500 in this analysis) number of abstracts is parsed and presented in the form of a table with paper titles and either a bar chart or word cloud of the most frequently occurring terms in these abstracts (Figure S2d).

In this way, iodine was shown to be often co-mentioned in abstracts with *thyroid*, *radiation* and *uptake,* among others. Now that the user has a quick overview of the current state of knowledge on this compound in this context, they can decide either to continue researching further leads within MetaboShiny, or move on to more focused literature research or follow-up experimentation, based on the potentially interesting leads described by the word cloud/ bar chart.

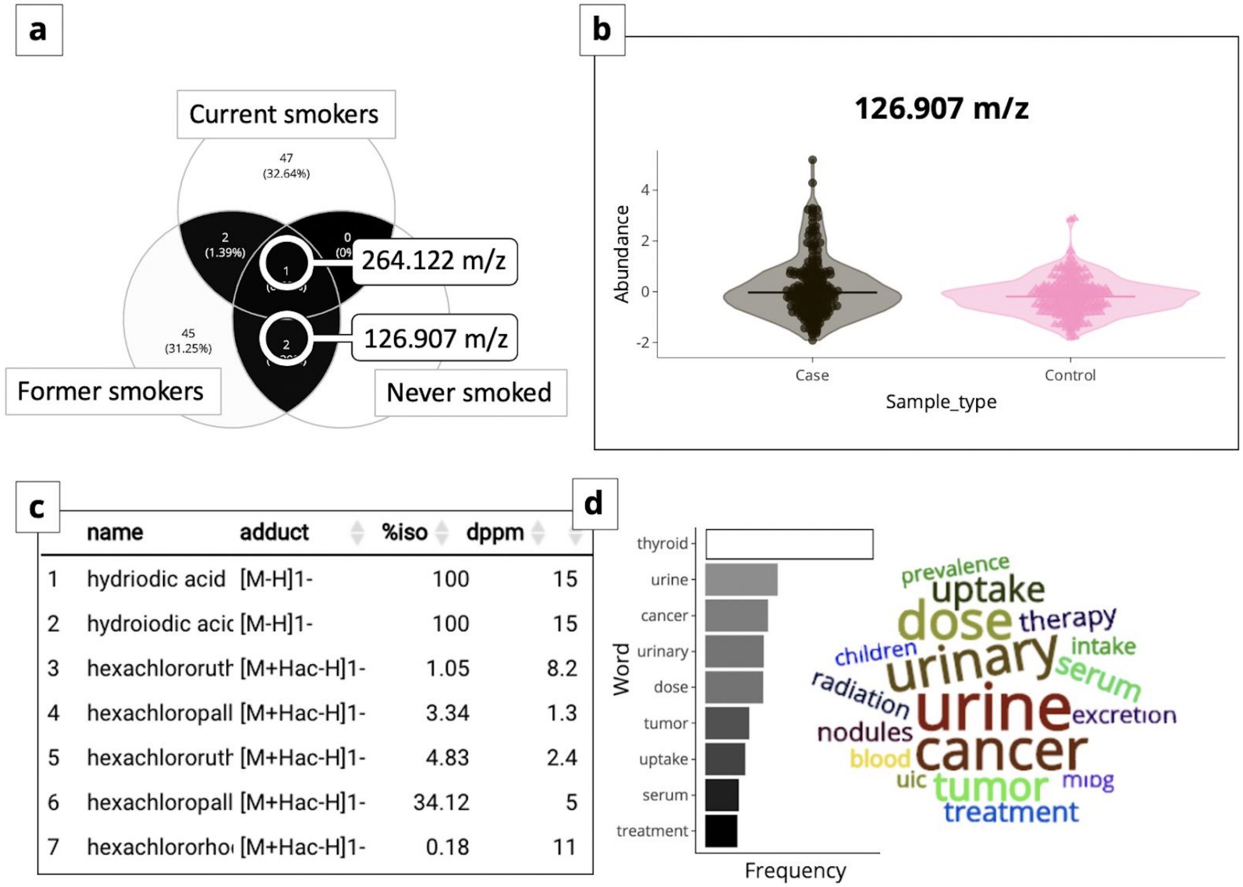


| | name | adduct | %iso | dppm |
|---|---|---|---|---|
| 1 | hydriodic acid | [M-H]1- | 100 | 15 |
| 2 | hydroiodic acic | [M-H]1- | 100 | 15 |
| 3 | hexachlororuth | [M+Hac-H]1- | 1.05 | 8.2 |
| 4 | hexachloropall | [M+Hac-H]1- | 3.34 | 1.3 |
| 5 | hexachlororuth | [M+Hac-H]1- | 4.83 | 2.4 |
| 6 | hexachloropall | [M+Hac-H]1- | 34.12 | 5 |
| 7 | hexachlororho | [M+Hac-H]1- | 0.18 | 11 |

**Figure S2:** Intersecting the top 50 predictive hits for smokers, nonsmokers and past smokers. (a) Venn diagram of overlap between the top 50 compounds included in final glmnet models for each subset. Metabolites of interest are circled.
(b) Differential expression of 127 m/z between lung cancer patients and population in nonsmokers. (c) Main search results in all databases. (d) Bar chart and word cloud of 500 PubMed abstracts matching the keywords "*iodine cancer urine*".

#### *Matching of a yet unannotated 561.3 m/z using Metaboshiny*

To identify the unknown metabolite with *m/z 561.3* the matching capability of MetaboShiny was leveraged. This compound was found to be significantly elevated in lung cancer patients by the original authors(Mathé *et al.*, 2014). While they established that the compound was conjugated with glucuronide (a process summarized in Figure S3a, generally to increase solubility and thus excretability(Sanchez and Kauffman, 2010)), its exact identity was not unraveled.

A new adduct, named *[M+GLUC±H],* was included in the search to represent glucuronidated compounds. To do this, knowledge on which chemical groups are necessary for glucuronidation was utilized to create rules in SMARTS format(*Table S4*)(Daylight Chemical Information, no date; Sanchez and Kauffman, 2010). Following the establishment of this new adduct, MetaboShiny's ability to flexibly calculate molecular adducts was used to implement it in the database(figure S3a).

All available databases returned their possible matches for *561.3 m/z*. When returning results, MetaboShiny offers the user a pie chart demonstrating from which databases the matches originate, and which adducts were mainly found as possible identities.

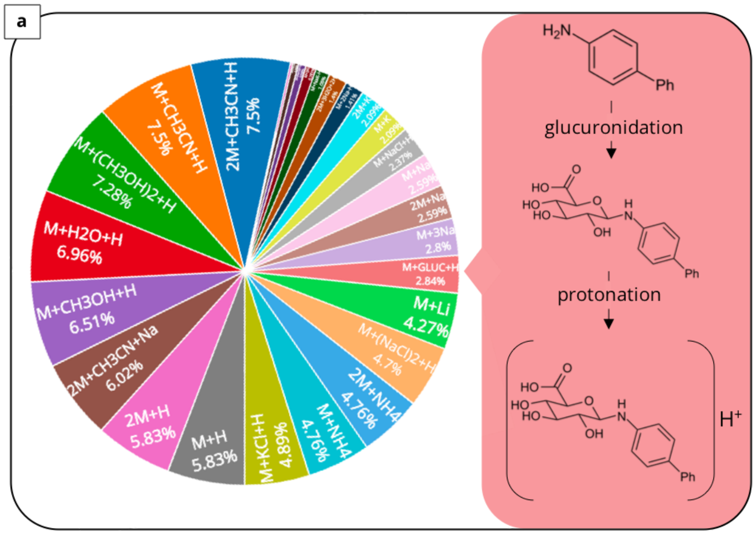


**b**

| name | adduct | %iso |
|---|---|---|
| **N-palmitoyl glutamine** | **M+GLUC+H** | **100** |
| L-759,633 | M+GLUC+H | 100 |
| L-759,656 | M+GLUC+H | 100 |
| sesquicannabigerol | M+GLUC+H | 100 |
| ethyl lauroyl arginate | M+GLUC+H | 100 |
| 5-Doxylstearate | M+GLUC+H | 100 |
| 16-Nitroxystearate | M+GLUC+H | 100 |
| BZLDGZBNMOECGO-UHFFFAOYSA-N | M+GLUC+H | 100 |

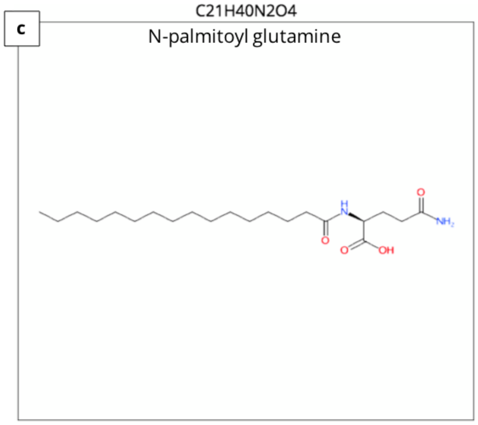


**d**

| pubmed ID | title |
|---|---|
| 28594046 | Balancing the intermolecular forces in peptide amphiphiles for controlling self-assembly transitions. |
| 28413374 | **Discrimination of pancreatic cancer and pancreatitis by LC-MS metabolomics.** |

**Figure S3:** Annotating 561.3 m/z, a glucuronidated compound. (a) Overview of all hits found for this m/z value shows that 2.9% of all hits are glucuronidated compounds. The right-hand box demonstrates glucuronidation. (b) Top hits of 16-database search for glucuronidated compounds. (c) Chemical structure of N-palmitoyl glutamine generated from SMILES structure description. (d) PubChem papers resulting from the integrated search. Used keyword was 'N-palmitoyl glutamic acid'.

Filtering out everything that was not a glucuronidated adduct through the interactive pie chart in *figure S3a* resulted in multiple main-isotope matches, which can be considered as possible matches for *561.3 m/z* (figure S3b). When creating new adducts one must consider if this is chemically valid. This is achieved by using the integrated molecular structure visualization (figure S3c) to estimate if the required groups for glucuronidation are present in the candidate molecule, enabled by the *rcdk* package(Guha and Cherto, 2017). Browsing through these results revealed N-palmitoyl glutamine.

The online PubChem search (figure S3d) furthermore uncovered a paper demonstrating the potential of a very closely related compound, N-palmitoyl glutamic acid, to differentiate pancreatic cancer from pancreatitis using LC-MS metabolomics(Lindahl *et al.*, 2017). Although this m/z value is found in a different setting, the fact that it is found in the context of LC-MS metabolomics in cancer detection makes this a strong lead to investigate further.

### Section S3. Comparison of Metaboshiny with existing tools

MetaboShiny performs multiple analysis steps and integrates many smaller tools into one toolbox. MetaboShiny should therefore be compared to toolboxes with a similar multi-level approach (Misra and Mohapatra, 2018). In this section we discuss some existing platforms and compare them to MetaboShiny (*Figure 2a*).

WebSpecMine is an online *Shiny*-based analysis tool. Key features are a direct connection to MetaboLights, data processing, normalization, statistics and annotation (Cardoso *et al.*, 2019). One limitation is that it only annotates LC-MS data. Furthermore, MetaboShiny offers a broader range of statistical analysis due to MetaboAnalyst integration, and the resulting plots are interactive.

MZmine is a downloadable open-source tool that allows for raw data processing, some statistical analyses and visualisation options (Olivon *et al.*, 2017). Although we have decided to not include raw data processing in our tool, we include more extensive statistical methods, including a machine learning model. Furthermore, MZmine is focussed on LC-MS data, and there is no guarantee that DI-MS data will be easy to import in this program.

Workflow4Metabolomics is a Galaxy-based tool going from raw data to statistics and tentative annotation(Guitton *et al.*, 2017). There is no direct support for DI-MS data, but the Galaxy platform is widely used and well known in the bioinformatics community, although it needs a user account. MetaboShiny provides more extensive statistical analyses and databases as well as interactivity of plots and tables.

Lastly, MetaboShiny has been built using the foundation of MetaboAnalyst with regard to normalization of the data and many of their statistical methods (Chong and Xia, 2018). MetaboShiny adds more interactivity, simultaneous analysis and annotation, additional machine learning and dimensionality reduction methods, and over 30 new metabolite databases.

| Tool name | DI-MS support | Machine learning | Interactive visualization | Can run locally | Open source | Annotation databases |
|---|---|---|---|---|---|---|
| workflow4metabolomics | ✗ | ✓ | ✗ | ~ | ✓ | **4** |
| MZmine | ~ | ~ | ~ | ✓ | ✓ | **9** |
| MetaboAnalyst | ~ | ✓ | ~ | ✓ | ✓ | **1** |
| WebSpecMine | ~ | ✓ | ✗ | ✗ | ✓ | **1** |
| **MetaboShiny** | ✓ | ✓ | ✓ | ✓ | ✓ | **35** |

**Table S1:** (a) Comparison of MetaboShiny to other multifunctional metabolomics tools (Guitton *et al.*, 2017; Olivon *et al.*, 2017; Chong *et al.*, 2018; Cardoso *et al.*, 2019). *Checkmark* (✓): full support, *tilde (~)*: partial support and *cross (x)*: no support.

### Table S2. Offline databases

*An overview of all the databases available for download in MetaboShiny. The advantage of being downloadable is that users can specify their own adducts to add to the database, allowing for extensive customization. It also allows for offline functionality and analysis. For each database, the name, logo (if available, otherwise other placeholder from the website), description and amount of unique structures (counted in # of SMILES) are listed in the table below. When taking into account the default adduct table (see Table S3) the final m/z count in the extended database including all isotopes and adducts is close to 200 million values.*

| Name | Logo | Official description | Unique structures |
|---|---|---|---|
| **HMDB** | | Metabolites commonly found in human biological samples. | 11 567 |
| **ChEBI** | | A broad database with known chemicals of biological interest. | 17 727 |
| **DimeDB** | | A direct infusion database of biologically relevant metabolite structures and annotations. | 1 4308 |
| **KEGG** | | Large pathway database with info on pathways in various organisms, involved enzymes, and connected disease phenotypes. | 8 957 |
| **MetaCyc** | | Large pathway database with over 10.000 available compounds. Spans several organisms. | 8 386 |
| **WikiPathways** | | Open source biological pathway database. Currently only partially available. Requires CHEBI to be built. | 880 |
| **SMPDB** | | Small molecule pathway database. Compounds overlap with HMDB. | 2 582 |
| **Wikidata** | | Central storage for the data of its Wikimedia sister projects including Wikipedia, Wikivoyage, Wikisource, and others. | 61 162 |

| | | | |
|---|---|---|---|
| **VMH** | | Virtual Metabolic Human (VMH) hosts ReconMap, an extensive network of human metabolism, and bacterial metabolites. | 3 149 |
| **ReSpect** | | RIKEN MSn spectral database for phytochemicals (ReSpect) is a collection of literature and in-house MSn spectra data for research on plant metabolomics. | 1 604 |
| **MassBank** | | This site presents the database of comprehensive, high-resolution mass spectra of metabolites. Supported by the JST-BIRD project, it offers various query methods for standard spectra from Keio Univ., RIKEN PSC, and others. | 6 822 |
| **MetaboLights** | | MetaboLights is a database for Metabolomics experiments and derived information. The database is cross-species, cross-technique and covers metabolite structures and their reference spectra as well as their biological roles, locations and concentrations, and experimental data from metabolic experiments. | 10 059 |
| **FooDB** | | FooDB is the world's largest and most comprehensive resource on food constituents, chemistry and biology. It provides information on both macronutrients and micronutrients, including many of the constituents that give foods their flavor, color, taste, texture and aroma. | 9 092 |
| **MaConDa** | | MaConDa currently contains ca. 200 contaminant records detected across several MS platforms. The majority of records include theoretical as well as experimental MS data. | 309 |
| **Blood Exposome Database** | | This new blood exposome database can be applied to prioritize literature-based chemical reviews, developing target assays in exposome research, identifying compounds in untargeted mass spectrometry and biological interpretation in metabolomics data. | 23 028 |
| **LipidMaps** | | The LIPID MAPS Structure Database (LMSD) is a relational database encompassing structures and annotations of biologically relevant lipids. | 7 964 |
| **Exposome Explorer** | | Exposome-Explorer is the first database dedicated to biomarkers of exposure to environmental risk factors for diseases. | 238 |
| **Toxin and Toxin Target Database(T3DB)** | | The Toxin and Toxin Target Database (T3DB), or, soon to be referred as, the Toxic Exposome Database, is a unique bioinformatics resource that combines detailed toxin data with comprehensive toxin target information. | 2 658 |
| **DrugBank** | | The DrugBank database is a unique bioinformatics and cheminformatics resource that combines detailed drug data with comprehensive drug target information. | 10 614 |

| | | | |
|---|---|---|---|
| **Phenol-Explorer** | Phenol-Explorer | Phenol-Explorer is the first comprehensive database on polyphenol content in foods. The database contains more than 35,000 content values for 500 different polyphenols in over 400 foods. | 737 |
| **PAMDB** | 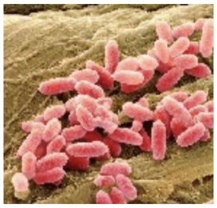 | The PAMDB is an expertly curated database containing extensive metabolomic data and metabolic pathway diagrams about Pseudomonas aeruginosa (reference strain PAO1). | 4 370 |
| **mVOC** | mVOC 2.0 | The mVOC 2.0 Database is based on extensive literature search for microbial volatile organic compounds (mVOCs) | 1 860 |
| **RMDB** | hmp | The Bovine Rumen Metabolome Database (RMDB) makes available tables containing the set of 246 ruminal fluid metabolites or metabolite species from the bovine ruminal fluid metabolome. | 335 |
| **BMDB** | hmp | The Bovine Metabolome Database (BMDB) is a freely available electronic database containing detailed information about small molecule metabolites found in beef and dairy cattle. | 7 975 |
| **STOFF-IDENT** | STOFF-IDENT | STOFF-IDENT is a database of water relevant substances collated from various sources within the STOFF-IDENT and FOR-IDENT projects, hosted by the Bavarian Environment Agency (Bayerisches Landesamt für Umwelt, LfU), the University of Applied Sciences Weihenstephan-Triesdorf (HSWT) and the Technical University of Munich (TUM). | 8 844 |
| **NANPDB** | NANPDB | To the best of our knowledge this is the largest database of natural products isolated from native organisms of Northern Africa. | 4 928 |
| **Unique:** | **~ 95.000** | **Unique m/z including adducts and isotopes** | **> 161 million** |

### Table S3. Online-only databases

*An overview of the databases available only online. There are two 'groups' of databases here, one being a direct m/z search engine in CEU Mass Mediator (which has access to KEGG, HMDB, LipidMaps, and most importantly, Metlin and MINE, generally difficult to obtain/access databases), and the other being databases that do not take into account adducts, and thus formula prediction needs to be done first through MetaboShiny. As in table S1, this table displays the name of the database, logo if available, official description and the amount of unique structures available for search queries. Due to being a direct m/z search, CEU Mass Mediator does not support the default adduct table and relies on a handful of custom adducts.*

| Name | Logo | Official description | Unique structures |
|---|---|---|---|
| **SUPER NATURAL II** | | A database of natural products. It contains 325,508 natural compounds (NCs), including information about the corresponding 2d structures, physicochemical properties, predicted toxicity class and potential vendors. | 325 508 |
| **PubChem** | | PubChem is the world's largest collection of freely accessible chemical information. | > 102 000 000 |
| **ChemSpider** | | A chemical structure database providing fast access to over 77 million structures, properties and associated information. | > 81 000 000 |
| **CEU Mass Mediator** | | CEU Mass Mediator is a tool for searching metabolites in different databases (Kegg, HMDB, LipidMaps, Metlin, MINE and an in-house library). | *unavailable* |
| **KNApSAcK** | | The purpose of the KNApSAcK Metabolomics is to search metabolites from MS peak, molecular weight and molecular formula, and species. | 51 179 |

### Table S4. Default adducts considered in database building

*An overview of the most recent table of considered/calculated adducts in database building and searching. Columns feature adduct name as shown in MetaboShiny, ion mode (either positive or negative - neutral adducts are considered by adding M- and M+ ions to the table), added charge(z), multiplier xM (for dimers and so on), Rem/AddAt which represent atoms added or deducted before multiplication, and AddEx/RemEx meaning atoms added or deducted after multiplication. Nelec is another measure of charge. The rule column represents a query in SMARTS that needs to return positive before each compound (in SMILES) is considered eligible for this adduct or deduct. This table was created by merging three published tables and adding our own modifications(MZedDB,(de Sain-van der Velden et al., 2017) and (Kind and Fiehn, 2007)).*

| Name | Ion_mode | z | xM | AddAt | RemAt | AddEx | RemEx | Nelec | Rule |
|---|---|---|---|---|---|---|---|---|---|
| [M+3Na]3+ | positive | 3 | 1 | | | Na3 | | -3 | Nacc>2 AND Nch=0 |
| [M+H+2Na]3+ | positive | 3 | 1 | | | Na2H1 | | -3 | Nacc>2 AND Nch=0 |
| [M+2H+Na]3+ | positive | 3 | 1 | | | Na1H2 | | -3 | Nacc>2 AND Nch=0 |
| [M+3H]3+ | positive | 3 | 1 | | | H3 | | -3 | Nacc>2 AND Nch=0 |
| [2M+3H2O+2H]2+ | positive | 2 | 2 | | | H8O3 | | -2 | Nacc>1 AND Nch=0 |
| [M+3ACN+2H]2+ | positive | 2 | 1 | | | C6H11N3 | | -2 | Nacc>1 AND Nch=0 |
| [M+2ACN+2H]2+ | positive | 2 | 1 | | | C4H8N2 | | -2 | Nacc>1 AND Nch=0 |
| [M+2Na]2+ | positive | 2 | 1 | | | Na2 | | -2 | Nacc>1 AND Nch=0 |
| [M+ACN+2H]2+ | positive | 2 | 1 | | | C2H5N1 | | -2 | Nacc>1 AND Nch=0 |
| [M+H+K]2+ | positive | 2 | 1 | | | K1H1 | | -2 | Nacc>1 AND Nch=0 |
| [M+H+Na]2+ | positive | 2 | 1 | | | Na1H1 | | -2 | Nacc>1 AND Nch=0 |
| [M+H+NH4]2+ | positive | 2 | 1 | | | N1H5 | | -2 | Nacc>1 AND Nch=0 |
| [M+2H]2+ | positive | 2 | 1 | | | H2 | | -2 | Nacc>1 AND Nch=0 |
| [M+2H-H2O-NH3]2+ | positive | 2 | 1 | | N1H2 | | O1 H1 | -2 | Nnhh>0 AND Noh>0 AND Nch=0 |
| [2M+ACN+Na]1+ | positive | 1 | 2 | | | C2H3Na1 N1 | | -1 | Nacc>0 AND Nch=0 |
| [2M+ACN+H]1+ | positive | 1 | 2 | | | C2H4N1 | | -1 | Nacc>0 AND Nch=0 |
| [2M+K]1+ | positive | 1 | 2 | | | K1 | | -1 | Nacc>0 AND Nch=0 |

| | | | | | | | | | |
|---|---|---|---|---|---|---|---|---|---|
| [2M+Na]1+ | positive | 1 | 2 | | | Na1 | | -1 | Nacc>0 AND Nch=0 |
| [2M+NH4]1+ | positive | 1 | 2 | | | N1H4 | | -1 | Nacc>0 AND Nch=0 |
| [2M+H]1+ | positive | 1 | 2 | | | H1 | | -1 | Nacc>0 AND Nch=0 |
| [M+IsoProp+Na+H]1+ | positive | 1 | 1 | | | C3H9O1 Na1 | | -1 | Nacc>0 AND Nch=0 |
| [M+2ACN+H]1+ | positive | 1 | 1 | | | C4H7N2 | | -1 | Nacc>0 AND Nch=0 |
| [M+DMSO+H]1+ | positive | 1 | 1 | | | C2H6OS1 | | -1 | Nacc>0 AND Nch=0 |
| [M+2K-H]1+ | positive | 1 | 1 | | | K2 | H1 | -1 | Ndon>0 AND Nch=0 |
| [M+ACN+Na]1+ | positive | 1 | 1 | | | C2H3Na1 N1 | | -1 | Nacc>0 AND Nch=0 |
| [M+IsoProp+H]1+ | positive | 1 | 1 | | | C3H9O1 Na1 | | -1 | Nacc>0 AND Nch=0 |
| [M+2Na-H]1+ | positive | 1 | 1 | | | Na2 | H1 | -1 | Ndon>0 AND Nch=0 |
| [M+ACN+H]1+ | positive | 1 | 1 | | | C2H4N1 | | -1 | Nacc>0 AND Nch=0 |
| [M+K]1+ | positive | 1 | 1 | | | K1 | | -1 | Nacc>0 AND Nch=0 |
| [M+H+CH3OH]1+ | positive | 1 | 1 | | | C1H5O1 | | -1 | Ndon>0 AND Nch=0 |
| [M+Na]1+ | positive | 1 | 1 | | | Na1 | | -1 | Nacc>0 AND Nch=0 |
| [M+NH4]1+ | positive | 1 | 1 | | | N1H4 | | -1 | Nacc>0 AND Nch=0 |
| [M+H]1+ | positive | 1 | 1 | | | H1 | | -1 | Nacc>0 AND Nch=0 |
| [M1+.]1+ | positive | 1 | 1 | | | | | 0 | Nch=1 |
| [M+H-NH3]1+ | positive | 1 | 1 | | N1H2 | | | -1 | Nnhh>0 AND Nch=0 |
| [M+H-H2O]1+ | positive | 1 | 1 | | O1H1 | | | -1 | Noh>0 AND Nch=0 |
| [M+H-FA]1+ | positive | 1 | 1 | | C1H1 O2 | | | -1 | Ncooh>0 AND Nch=0 |
| M | positive | 1 | 1 | | | | | 0 | Nch=0 |
| [M+GLUC+H]1+ | positive | 1 | 1 | | | C6H10O 6 | H1 | -1 | Ngluc>0 AND Nch=0 |
| [3M-H]1- | negative | -1 | 3 | | | | H1 | 1 | Ndon>0 AND Nch=0 |
| [2M+Hac-H]1- | negative | -1 | 2 | | | C2H3O2 | | 1 | Ndon>0 AND Nch=0 |

| | | | | | | | | | |
|---|---|---|---|---|---|---|---|---|---|
| [2M+FA-H]1- | negative | -1 | 2 | | | C1H1O2 | | 1 | Ndon>0 AND Nch=0 |
| [2M+Na-2H]1- | negative | -1 | 2 | | | Na1 | H2 | 1 | Ndon>1 AND Nacc>0 AND Nch=0 |
| [2M-H]1- | negative | -1 | 2 | | | | H1 | 1 | Ndon>0 AND Nch=0 |
| [M+TFA-H]1- | negative | -1 | 1 | | | C2O2F3 | | 1 | Ndon>0 AND Nch=0 |
| [M+Br]1- | negative | -1 | 1 | | | Br1 | | 1 | Nacc>0 AND Nch=0 |
| [M+Hac-H]1- | negative | -1 | 1 | | | C2H3O2 | | 1 | Ndon>0 AND Nch=0 |
| [M+FA-H]1- | negative | -1 | 1 | | | C1H1O2 | | 1 | Ndon>0 AND Nch=0 |
| [M+K-2H]1- | negative | -1 | 1 | | | K1 | H2 | 1 | Ndon>1 AND Nacc>0 AND Nch=0 |
| [M+Cl]1- | negative | -1 | 1 | | | Cl1 | | 1 | Nacc>0 AND Nch=0 |
| [M+Na-2H]1- | negative | -1 | 1 | | | Na1 | H2 | 1 | Ndon>1 AND Nacc>0 AND Nch=0 |
| [M1-.]1- | negative | -1 | 1 | | | | | 0 | Nch=-1 |
| [M-H]1- | negative | -1 | 1 | | | | H1 | 1 | Ndon>0 AND Nch=0 |
| [M-2H]2- | negative | -2 | 1 | | | | H2 | 2 | Ndon>1 AND Nch=0 |
| [M-3H]3- | negative | -3 | 1 | | | | H3 | 3 | Ndon>2 AND Nch=0 |
| [M-H+GLUC]1- | negative | -1 | 1 | | | C6H9O6 | H2 | 1 | Ngluc>0 AND Nch=0 |

### Table S5. Rules for adduct formation

*Companion table to the adduct table S3. This table is used to define the adduct rules used in the default adduct table. Each rule is given a name for easier referencing, a descriptor of what the rule was made for, and, most importantly, a SMARTS query (except for the ‘charge/Nch’ rule, which is an exception and derived from the base molecule charge). Users can add their own here. We devised these rules by finding matching SMARTS for the rules introduced by MzedDB using the SMARTS Theory Manual (Daylight Chemical Information, no date).*

| Short name | Description | SMARTS |
|---|---|---|
| Nch | number of charges in M. | *Get charge from SMILES* |
| Nacc | number of H-bond acceptor in M. | [!$([#6,F,Cl,Br,I,o,s,nX3,#7v5,#15v5,#16v4,#16v6,*+1,*+2,*+3])] |
| Ndon | number of H-bond donor in M. | [!$([#6,H0,-,-2,-3])] |
| Noh | number of -OH groups in M. | [OX2H] |
| Ncooh | number of -COOH groups in M. | [CX3](=O)[OX2H1] |
| Ncoo | number of -COO- groups in M. | [CX3](=O)[O-] |
| Nnhh | number of -NH2 groups in M. | [NX3;H2,H1;!$(NC=O)] |
| Naci | number of acidic H in M. | [H+] |
| Nbas | number of basic O- in M. | [O-] |
| Ngluc | number of groups in M that can be glucuronidated | [$([#6][OX2H]),$([OX2H][CX3]=[OX1]),$([NX3,NX4+][CX3](=[OX1])[OX2H,OX1-]),$([NX2:2][OH1:3]),$([NX3]-[CX3](=[OX1])[OH]),$([NX3][CX3](=[OX1])[#6]),$([N;!H0;$(N-c);!$(N-[!#6;!#1]);!$(N-C=[O,N,S])]),$([N;!H0;!$(N-c);$(N-C);!$(N-[!#6;!#1]);!$(N-C=[O,N,S])]),$([#16X2H]),$([SX4](=[OX1])(=[OX1])([!O])[NX3]),$([SX4+2]([OX1-])([OX1-])([!O])[NX3]),$([CX3](=O)[CH2][CX3](=O))] |

### Table S6. Example metadata table

*Example of metadata table. Users can also use the MetaboLights default template(Daylight Chemical Information, no date; Kale et al., 2016). If using this format, the 'sample', and 'individual' columns are required, the others are freeform and users can add as much as they please. In this case, the user could load the data into MetaboShiny and do statistics on group, sex, or do time series analysis on these features as each patient has multiple samples in the database.*

| sample | individual | date | sex | group |
|---|---|---|---|---|
| **PAT1_1** | PAT1 | 4-Jun-19 | Male | Treatment A |
| **PAT1_2** | PAT1 | 14-Jun-19 | Male | Treatment A |
| **PAT1_3** | PAT1 | 24-Jun-19 | Male | Treatment A |
| **PAT2_1** | PAT2 | 4-Jun-19 | Female | Treatment B |
| **PAT2_2** | PAT2 | 14-Jun-19 | Female | Treatment B |
| **PAT2_3** | PAT2 | 24-Jun-19 | Female | Treatment B |
| **PAT3_1** | PAT3 | 4-Jun-19 | Male | Treatment C |
| **PAT3_2** | PAT3 | 14-Jun-19 | Male | Treatment C |
| **PAT3_3** | PAT3 | 24-Jun-19 | Male | Treatment C |

### Table S7. Example peak table

*Example of peak table. Separate peak tables need to be input for each ion mode (positive and negative). Users can also use the MetaboLights default template (Daylight Chemical Information, no date; Kale et al., 2016). Columns after the first column feature m/z values, and rows represent samples. Intensities can be normalized or non-normalized - however it is important to not normalize the data again if that has already been done.*

| sample | 71 | 72 | 73 | 74 |
|---|---|---|---|---|
| **PAT1_1** | 12506 | 234905 | 23490 | 349 |
| **PAT1_2** | 23940 | 33494 | 29385 | 29485 |
| **PAT1_3** | 28305 | 2930 | 923 | 20395 |
| **PAT2_1** | *NA* | 92843 | 9203 | 2039 |
| **PAT2_2** | 3849 | *NA* | 238940 | 2189 |
| **PAT2_3** | 23894 | 9023 | 902 | 23890 |
| **PAT3_1** | 5863 | 86739 | 9283 | 293 |
| **PAT3_2** | 82734 | 29385 | 2389 | 67983 |
| **PAT3_3** | 1283959 | 38495 | 1029 | 23895 |

### Section S3. Supplemental references


Allen, G. and Owens, M. (2010) “SQL for SQLite,” in Allen, G. and Owens, M. (eds.) *The Definitive Guide to SQLite*. Berkeley, CA: Apress, pp. 47–86. doi: 10.1007/978-1-4302-3226-1_3.

Anderson, E., Veith, G. D. and Weininger, D. (1987) “SMILES: A line notation and computerized interpreter for chemical structures. Duluth, MN: US EPA,” *Environmental Research Laboratory-Duluth. Report No. EPA/600/M-87/021*.

Ayers, M. (2012) “ChemSpider: The Free Chemical Database2012312ChemSpider: The Free Chemical Database. URL: www.chemspider.com: Royal Society of Chemistry Last visited April 2012. Gratis,” *Reference Reviews*, pp. 45–46. doi: 10.1108/09504121211271059.

Barupal, D. K. and Fiehn, O. (2019) “Generating the Blood Exposome Database Using a Comprehensive Text Mining and Database Fusion Approach,” *Environmental health perspectives*, 127(9), p. 97008. doi: 10.1289/EHP4713.

Cardoso, S. *et al.* (2019) “WebSpecmine: A Website for Metabolomics Data Analysis and Mining,” *Metabolites*, 9(10). doi: 10.3390/metabo9100237.

Caspi, R. *et al.* (2020) “The MetaCyc database of metabolic pathways and enzymes - a 2019 update,” *Nucleic acids research*, 48(D1), pp. D445–D453. doi: 10.1093/nar/gkz862.

Chong, J. *et al.* (2018) “MetaboAnalyst 4.0: towards more transparent and integrative metabolomics analysis,” *Nucleic acids research*, 46(W1), pp. W486–W494. doi: 10.1093/nar/gky310.

Chong, J. and Xia, J. (2018) “MetaboAnalystR: an R package for flexible and reproducible analysis of metabolomics data,” *Bioinformatics* , 34(24), pp. 4313–4314. doi: 10.1093/bioinformatics/bty528.

Daylight Chemical Information (no date) “SMARTS Theory Manual.” Available at: https://www.daylight.com/dayhtml/doc/theory/theory.smarts.html.

Degtyarenko, K. *et al.* (2008) “ChEBI: a database and ontology for chemical entities of biological interest,” *Nucleic acids research*, 36(Database issue), pp. D344–50. doi: 10.1093/nar/gkm791.

Dunkel, M. *et al.* (2006) “SuperNatural: a searchable database of available natural compounds,” *Nucleic acids research*, 34(Database issue), pp. D678–83. doi: 10.1093/nar/gkj132.

Gil-de-la-Fuente, A. *et al.* (2019) “CEU Mass Mediator 3.0: A Metabolite Annotation Tool,” *Journal of proteome research*, 18(2), pp. 797–802. doi: 10.1021/acs.jproteome.8b00720.

Guha, R. and Cherto, M. R. (2017) “rcdk: Integrating the CDK with R.” Available at: http://www.idg.pl/mirrors/CRAN/web/packages/rcdk/vignettes/rcdk.pdf.

Guitton, Y. *et al.* (2017) “Create, run, share, publish, and reference your LC-MS, FIA-MS, GC-MS, and NMR data analysis workflows with the Workflow4Metabolomics 3.0 Galaxy online infrastructure for metabolomics,” *The international journal of biochemistry & cell biology*, 93, pp. 89–101. doi: 10.1016/j.biocel.2017.07.002.

Guo, A. C. *et al.* (2012) “ECMDB: The E. coli Metabolome Database,” *Nucleic Acids Research*, pp. D625–D630. doi: 10.1093/nar/gks992.

Horai, H. *et al.* (2010) “MassBank: a public repository for sharing mass spectral data for life sciences,”

*Journal of Mass Spectrometry*, pp. 703–714. doi: 10.1002/jms.1777.

Huang, W. *et al.* (2018) "PAMDB: a comprehensive Pseudomonas aeruginosa metabolome database," *Nucleic acids research*, 46(D1), pp. D575–D580. doi: 10.1093/nar/gkx1061.

Ihlenfeldt, W.-D. (2018) "PubChem," *Applied Chemoinformatics*, pp. 245–258. doi: 10.1002/9783527806539.ch6e.

Jewison, T. *et al.* (2014) "SMPDB 2.0: big improvements to the Small Molecule Pathway Database," *Nucleic acids research*, 42(Database issue), pp. D478–84. doi: 10.1093/nar/gkt1067.

Johnson, W. E., Li, C. and Rabinovic, A. (2007) "Adjusting batch effects in microarray expression data using empirical Bayes methods," *Biostatistics* , 8(1), pp. 118–127. doi: 10.1093/biostatistics/kxj037.

Kale, N. S. *et al.* (2016) "MetaboLights: An Open-Access Database Repository for Metabolomics Data," *Current protocols in bioinformatics / editoral board, Andreas D. Baxevanis ... [et al.]*. Wiley Online Library, 53(1), pp. 14–13. Available at: https://currentprotocols.onlinelibrary.wiley.com/doi/abs/10.1002/0471250953.bi1413s53.

Kanehisa, M. and Goto, S. (2000) "KEGG: kyoto encyclopedia of genes and genomes," *Nucleic acids research*, 28(1), pp. 27–30. Available at: https://www.ncbi.nlm.nih.gov/pubmed/10592173.

Kind, T. and Fiehn, O. (2007) "Seven Golden Rules for heuristic filtering of molecular formulas obtained by accurate mass spectrometry," *BMC bioinformatics*, 8, p. 105. doi: 10.1186/1471-2105-8-105.

Kuhn, M. (2008) "Building predictive models in R using the caret package," *Journal of statistical software*. math.chalmers.se. Available at: http://www.math.chalmers.se/Stat/Grundutb/GU/MSA220/S18/caret-JSS.pdf.

Lemfack, M. C. *et al.* (2018) "mVOC 2.0: a database of microbial volatiles," *Nucleic acids research*, 46(D1), pp. D1261–D1265. doi: 10.1093/nar/gkx1016.

Lindahl, A. *et al.* (2017) "Discrimination of pancreatic cancer and pancreatitis by LC-MS metabolomics," *Metabolomics: Official journal of the Metabolomic Society*, 13(5), p. 61. doi: 10.1007/s11306-017-1199-6.

Mathé, E. A. *et al.* (2014) "Noninvasive urinary metabolomic profiling identifies diagnostic and prognostic markers in lung cancer," *Cancer research*, 74(12), pp. 3259–3270. doi: 10.1158/0008-5472.CAN-14-0109.

Misra, B. B. and Mohapatra, S. (2018) "Tools and resources for metabolomics research community: A 2017–2018 update," *Electrophoresis*. Wiley Online Library. Available at: https://onlinelibrary.wiley.com/doi/abs/10.1002/elps.201800428.

Nakamura, Y. *et al.* (2014) "KNApSAcK Metabolite Activity Database for retrieving the relationships between metabolites and biological activities," *Plant & cell physiology*, 55(1), p. e7. doi: 10.1093/pcp/pct176.

Neveu, V. *et al.* (2020) "Exposome-Explorer 2.0: an update incorporating candidate dietary biomarkers and dietary associations with cancer risk," *Nucleic acids research*, 48(D1), pp. D908–D912. doi: 10.1093/nar/gkz1009.

Noronha, A. *et al.* (2019) "The Virtual Metabolic Human database: integrating human and gut microbiome metabolism with nutrition and disease," *Nucleic acids research*, 47(D1), pp. D614–D624. doi: 10.1093/nar/gky992.

Ntie-Kang, F. *et al.* (2017) "NANPDB: A Resource for Natural Products from Northern African Sources," *Journal of natural products*, 80(7), pp. 2067–2076. doi: 10.1021/acs.jnatprod.7b00283.

Olivon, F. *et al.* (2017) "MZmine 2 Data-Preprocessing To Enhance Molecular Networking Reliability," *Analytical Chemistry*, pp. 7836–7840. doi: 10.1021/acs.analchem.7b01563.

O'Shea, K. *et al.* (no date) "DIMEdb: an integrated database and web service for metabolite identification in direct infusion mass spectrometery." doi: 10.1101/291799.

Rothwell, J. A. *et al.* (2013) "Phenol-Explorer 3.0: a major update of the Phenol-Explorer database to incorporate data on the effects of food processing on polyphenol content," *Database*, pp. bat070–bat070. doi: 10.1093/database/bat070.

de Sain-van der Velden, M. G. M. *et al.* (2017) "Quantification of metabolites in dried blood spots by direct infusion high resolution mass spectrometry," *Analytica chimica acta*, 979, pp. 45–50. doi: 10.1016/j.aca.2017.04.038.

Sanchez, R. I. and Kauffman, F. C. (2010) "Regulation of Xenobiotic Metabolism in the Liver," *Comprehensive Toxicology*, pp. 109–128. doi: 10.1016/b978-0-08-046884-6.01005-8.

Sawada, Y. *et al.* (2012) "RIKEN tandem mass spectral database (ReSpect) for phytochemicals: A plant-specific MS/MS-based data resource and database," *Phytochemistry*, pp. 38–45. doi: 10.1016/j.phytochem.2012.07.007.

Sievert, C. *et al.* (2016) "plotly: Create Interactive Web Graphics via 'plotly. js,'" *R package version*.

Slenter, D. N. *et al.* (2018) "WikiPathways: a multifaceted pathway database bridging metabolomics to other omics research," *Nucleic acids research*, 46(D1), pp. D661–D667. doi: 10.1093/nar/gkx1064.

Stekhoven, D. J. (2015) "missForest: Nonparametric missing value imputation using random forest," *Astrophysics Source Code Library*. Available at: https://ui.adsabs.harvard.edu/#abs/2015ascl.soft05011S.

Subramaniam, S. and Fahy, E. (2007) "LipidMaps Core Update," *Nature Precedings*. doi: 10.1038/npre.2007.23.1.

Turki, H. *et al.* (2019) "Wikidata: A large-scale collaborative ontological medical database," *Journal of biomedical informatics*, 99, p. 103292. doi: 10.1016/j.jbi.2019.103292.

Weber, R. J. M. *et al.* (2012) "MaConDa: a publicly accessible mass spectrometry contaminants database," *Bioinformatics* , 28(21), pp. 2856–2857. doi: 10.1093/bioinformatics/bts527.

Wehrens, R. *et al.* (2016) "Improved batch correction in untargeted MS-based metabolomics," *Metabolomics: Official journal of the Metabolomic Society*, 12, p. 88. doi: 10.1007/s11306-016-1015-8.

Wei, R. *et al.* (2018) "Missing Value Imputation Approach for Mass Spectrometry-based Metabolomics Data," *Scientific reports*, 8(1), p. 663. doi: 10.1038/s41598-017-19120-0.

Wickham, H. (2011) "ggplot2," *Wiley Interdisciplinary Reviews: Computational Statistics*, 3(2), pp. 180–185. doi: 10.1002/wics.147.

Wishart, D. *et al.* (2015) "T3DB: the toxic exposome database," *Nucleic acids research*, 43(Database issue), pp. D928–34. doi: 10.1093/nar/gku1004.

Wishart, D. S., Feunang, Y. D., *et al.* (2018) "DrugBank 5.0: a major update to the DrugBank database for 2018," *Nucleic Acids Research*, pp. D1074–D1082. doi: 10.1093/nar/gkx1037.

Wishart, D. S., Feunang, Y. D., *et al.* (2018) “HMDB 4.0: the human metabolome database for 2018,” *Nucleic acids research*, 46(D1), pp. D608–D617. doi: 10.1093/nar/gkx1089.

Xie, Y. (2017) “DT: a wrapper of the JavaScript library ‘DataTables’. 2016,” *R package version 0. 2*.
